# Angpt1 binding to Tie1 regulates the signaling required for lymphatic vessel development in zebrafish

**DOI:** 10.1101/2023.08.16.553481

**Authors:** Nanami Morooka, Ning Gui, Koji Ando, Keisuke Sako, Moe Fukumoto, Melina Hußmann, Stefan Schulte-Merker, Naoki Mochizuki, Hiroyuki Nakajima

**Affiliations:** Department of Cell Biology, National Cerebral and Cardiovascular Center Research Institute, Suita, Osaka 564-8565, Japan; Department of Medical Physiology, Hamamatsu University School of Medicine, Hamamatsu, Shizuoka 431-3192, Japan; Department of Cardiac Regeneration Biology, National Cerebral and Cardiovascular Center Research Institute, Suita, Osaka 564-8565, Japan; Institute of Cardiovascular Organogenesis and Regeneration, Faculty of Medicine, WWU Münster, 48149 Münster, Germany

**Keywords:** Lymphangiogenesis, Tie1, Zebrafish, Live imaging

## Abstract

Development of the vascular system is regulated by multiple signaling pathways mediated by receptor tyrosine kinases (RTKs). Among them, Angiopoietin (Ang)/Tie signaling regulates lymphatic and blood vessel development in mammals. Of the two Tie receptors, Tie2 is well known as a key mediator of Ang/Tie signaling, but unexpectedly, recent studies reveal that the *Tie2* locus has been lost in many vertebrate species, while the *Tie1* gene is more commonly present. However, Tie1-driven signaling pathways, including ligands and cellular functions, are not well understood. Here, we performed comprehensive mutant analyses of Angiopoietins and Tie receptors in zebrafish and found that only *angpt1* and *tie1* mutants show defects in trunk lymphatic vessel development. Among zebrafish Angiopoietins, only Angpt1 binds to Tie1 as a ligand. We indirectly monitored Ang1/Tie1 signaling and detected Tie1 activation in sprouting endothelial cells (ECs), where Tie1 inhibits nuclear import of EGFP-Foxo1a. Angpt1/Tie1 signaling functions in EC migration, proliferation, and lymphatic specification during early lymphangiogenesis, at least in part by modulating Vegfc/Vegfr3 signaling. Thus, we show Angpt1/Tie1 signaling to constitute an essential signaling pathway for lymphatic development in zebrafish.

**Brief Summary Statement:** Zebrafish Angpt1/Tie1 signaling is characterized as an essential signaling pathway for trunk lymphatic development, with Tie1 regulating Foxo1 localization and modulating Vegfc/Vegfr3 signaling.

## INTRODUCTION

Lymphatic vessels, together with heart and blood vessels, constitute the vascular system and play pivotal roles in the maintenance of fluid balance, fat absorption, and immunosurveillance (Alitalo, 2011; Donnan et al., 2021; Escobedo and Oliver, 2016; Francois et al., 2021; Petrova and Koh, 2020). During development, most lymphatic vessels derive from preexisting blood vessels. At the onset of lymphangiogenesis, a subpopulation of venous endothelial cells (ECs) within the large veins, such as the cardinal vein (CV) in mice and the posterior cardinal vein (PCV) in zebrafish, express the transcription factor Prox1, a well-known marker of lymphatic ECs (LECs), and thereby commit to a lymphatic endothelial cell fate (Koltowska et al., 2015; Nicenboim et al., 2015; Wigle and Oliver, 1999). These Prox1^+^ LEC progenitors sprout from the vein and undergo migration, proliferation, and differentiation, eventually giving rise to the primary lymphatic vessels (Escobedo and Oliver, 2016; Suarez and Schulte-Merker, 2021). An endothelial-specific receptor tyrosine kinase (RTK) Vegfr3 (also known as Flt4) and its ligand Vegfc play a central role in lymphangiogenesis both in zebrafish and mice (Hogan et al., 2009; Jeltsch et al., 2013; Karkkainen et al., 2004; Le Guen et al., 2014; Tammela and Alitalo, 2010; Villefranc et al., 2013). Vegfc/Vegfr3 signaling is essential for lymphatic differentiation and the initial sprouting of lymphatic progenitors from the vein (Karkkainen et al., 2004; Koltowska et al., 2015; Shin et al., 2016; Srinivasan et al., 2014; Tammela and Alitalo, 2010). This process of primary lymphatic vasculature formation is conserved from fish to mammals (Hagerling et al., 2013; Mauri et al., 2018).

The Angiopoietin/Tie system is another endothelial-specific RTK signaling important for mammalian cardiovascular development. Of the two TIE receptors, only TIE2 has been reported to bind to its ligands, Angiopoietins, in mammals (Augustin et al., 2009; Eklund and Saharinen, 2013). Therefore, TIE2 is thought to be a key mediator of Ang/Tie signaling. Among Angiopoietins, Angiopoietin 1 (ANG1) is a constitutive agonist for TIE2 phosphorylation that is crucial for blood vessel integrity (Fukuhara et al., 2008; Puri et al., 1995; Saharinen et al., 2008), whereas ANG2 is a context-dependent TIE2 agonist or antagonist (Daly et al., 2006; Kim et al., 2016; Maisonpierre et al., 1997). During lymphatic development, ANG2 acts as a TIE2 agonist (Gale et al., 2002). In contrast, mammalian TIE1 is an orphan receptor and functions by forming a heterodimer with TIE2, thereby acting as a context-dependent modulator of TIE2 activity (Korhonen et al., 2016; Saharinen et al., 2005; Savant et al., 2015; Seegar et al., 2010; Yuan et al., 2007). However, it is noteworthy that some studies show a unique role of TIE1 in lymphatic vessel formation and maturation. Hypomorphic mice with reduced expression of *Tie1* show lymphatic vascular abnormalities and prominent edema (D’Amico et al., 2010; Qu et al., 2010). Furthermore, postnatal deletion of *Tie1* causes abnormalities in the lymphatic capillary network, whereas deletion of *Tie2* does not (Korhonen et al., 2022; Shen et al., 2014). These reports highlight the crucial role of TIE1 as the dominant TIE receptor required for lymphatic vessel formation. However, how Tie1 regulates lymphangiogenesis is not fully understood.

Recent studies unexpectedly reveal that, of the two Tie receptors, the *tie1* gene is more commonly present in vertebrates, but the *tie2* gene is absent in most of the ray-finned fish, including Medaka (Oryzias latipes) (Gjini et al., 2011; Jiang et al., 2020). Although the *tie2* gene is present in zebrafish, no obvious phenotypes have been reported in *tie2* mutants (Gjini et al., 2011; Jiang et al., 2020). In contrast, zebrafish *tie1* mutants die primarily due to cardiac defects (Carlantoni et al., 2021). These mutants exhibit vascular defects such as reduced EC numbers, reduced blood flow, impaired caudal vein plexus formation, and impaired lymphatic development (Carlantoni et al., 2021; Hußmann et al., 2023). Therefore, Tie1 may be responsible for Ang/Tie signaling in cardiovascular development in most fish species that lack functional Tie2. However, there have been no reports on the roles of Angiopoietins in these species. In addition, Tie1-driven signaling pathways, including ligands, downstream signaling, and cellular functions, remain unclear in any animal models.

In this study, we performed comprehensive functional analyses of Angiopoietins and Tie receptors in zebrafish. Our *in vitro* binding assay shows that, among Angiopoietins, only Angpt1 (zebrafish ortholog of Ang1) binds to Tie1 as a ligand. Similarly, *angpt1* mutants, unlike *angpt2a* and *angpt2b* mutants, phenocopy *tie1* mutants, which show defects in lymphatic vessel development. Angpt1/Tie1 signaling thus functions in early lymphangiogenesis in the trunk, where Tie1 negatively regulates nuclear translocation of Foxo1a downstream of PI3K and plays an essential role in the full activation of Vegfc/Vegfr3 signaling. We present evidence that Tie1 functions as a receptor for Angiopoietin and constitutes an essential signaling pathway for lymphatic development in zebrafish.

## RESULTS

### Angpt1 and Tie1 are essential for the development of lymphatic vessels in the trunk of zebrafish

In this study, we investigate the roles of Ang/Tie signaling in zebrafish by analyzing *angiopoietin* genes (*angpt1*, *angpt2a*, and *angpt2b*) and *tie* genes (*tie1* and *tie2*). To accomplish this, we generated mutants of *angpt1*, *angpt2a*, *angpt2b* as well as *tie1* by TALEN (Fig. S1A) and compared possible phenotypes with an established *tie2* mutant (Gjini et al., 2011). As reported previously (Gjini et al., 2011; Jiang et al., 2020), homozygous *tie2* mutants did not show any obvious phenotypes in lymphatic or blood vessel development (Fig. 1A,C). In contrast, homozygous *tie1* mutant larvae showed a loss of lymphatic structures in the trunk (Fig. 1A), while they did form trunk blood vessels (Fig. S1B). In these mutants, formation of the thoracic duct (TD), the first functional lymphatic vessel along the dorsal aorta (DA), was severely disrupted (Fig. 1A,B). In addition, *tie1* mutants exhibited cardiac edema from 2.5 dpf (Carlantoni et al., 2021) and severe edema around eyes and intestine from 4 dpf (Fig. 1C,D and Fig. S1C). Finally, all homozygous *tie1* mutants died before reaching adulthood.

**Figure 1.**
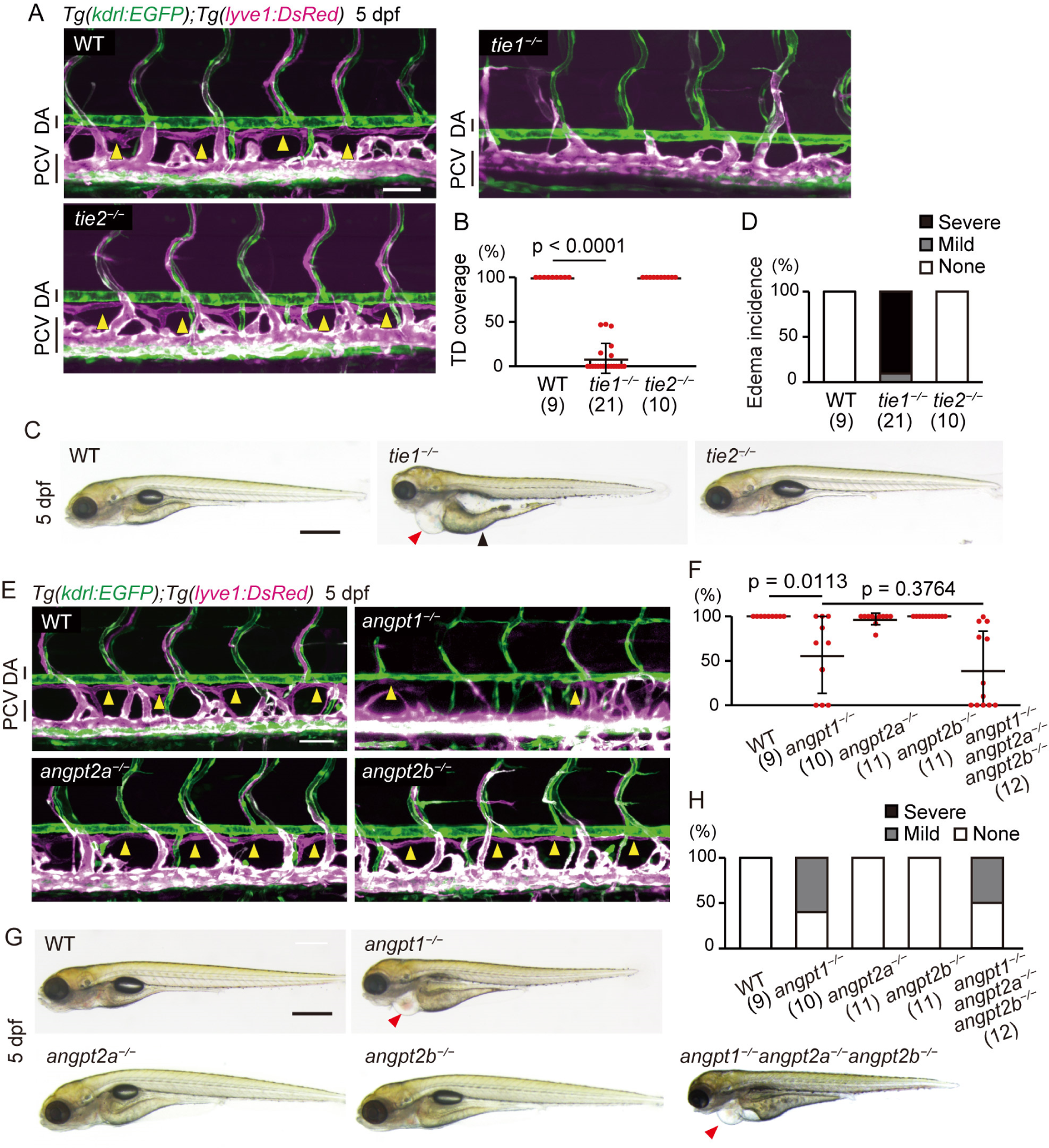
Angpt1 and Tie1 are essential for trunk lymphangiogenesis in zebrafish. (**A**) Representative confocal images of the trunk of *Tg(kdrl:EGFP);Tg(lyve1:DsRed)* wild-type (WT), *tie1^−/−^*, and *tie2^−/−^* larvae (5 dpf). These Tg larvae express EGFP (green) in all endothelial cells (ECs) and DsRed (magenta) in venous and lymphatic ECs. Formation of the thoracic duct (TD, yellow arrowheads in WT and *tie2*^−/−^ larvae) was impaired in *tie1^−/−^* larvae. Lateral views, anterior to the left. (**B**) Percentage of TD coverage at 5 dpf. Data are mean ± s.d. (WT, n = 9 larvae; *tie1^−/−^*, n = 21 larvae; *tie2^−/−^*, n = 10 larvae). (**C**) Overall morphology of WT, *tie1^−/−^*, and *tie2^−/−^* larvae (5 dpf). Red and black arrowheads point to edema around the heart and intestine, respectively. (**D**) Percentage of edema incidence at 5 dpf. Data are mean ± s.d. (WT, n = 9 larvae; *tie1^−/−^*, n = 21 larvae; *tie2^−/−^*, n = 10 larvae). Typical pictures of each category are shown in Fig. S1C. (**E**) Trunk of *Tg(kdrl:EGFP);Tg(lyve1:DsRed)* WT, *angpt1^−/−^*, *angpt2a^−/−^*, and *angpt2b^−/−^* larvae (5 dpf). Formation of the TD (yellow arrowheads in WT, *angpt2a^−/−^*, and *angpt2b^−/−^*larvae) was impaired in the *angpt1^−/−^* larvae. (**F**) Percentage of TD coverage at 5 dpf. Data are mean ± s.d. (WT, n = 9 larvae; *angpt1^−/−^*, n = 10 larvae; *angpt2a^−/−^*, n = 11 larvae; *angpt2b^−/−^*, n = 11 larvae; *angpt1^−/−^angpt2a^−/−^angpt2b^−/−^*, n = 12 larvae). (**G**) Overall morphology of WT, *angpt1^−/−^*, *angpt2a^−/−^*, and *angpt2b^−/−^*, and *angpt1^−/−^angpt2a^−/−^angpt2b^−/−^* triple mutant larvae (5 dpf). Arrowheads point to edema around the heart. (**H**) Percentage of edema incidence at 5 dpf as in (D). Data are mean ± s.d. (WT, n = 9 larvae; *angpt1^−/−^*, n = 10 larvae; *angpt2a^−/−^*, n = 11 larvae; *angpt2b^−/−^*, n = 11 larvae; *angpt1^−/−^angpt2a^−/−^angpt2b^−/−^*, n = 12 larvae). Scale bars: 50 μm (A,E), 500 μm (C,G). DA, dorsal aorta; PCV, posterior cardinal vein.

The function of mammalian TIE1 is considered to depend upon its binding to TIE2 (Saharinen et al., 2005; Yuan et al., 2007). To test whether the function of zebrafish Tie1 depends on Tie2, we analyzed *tie1*-*tie2* double mutants. Impaired lymphatic development and severe edema formation observed in *tie1* mutants were not rescued in *tie1*-*tie2* double mutants (Fig. S1D-G), indicating that the phenotypes of *tie1* mutants do not depend on the presence of Tie2 in zebrafish. Therefore, zebrafish Tie1 (hereafter referred to as zTie1) regulates lymphatic development independently of zTie2.

Among *angiopoietin* mutants, *angpt1* mutants exhibited cardiac edema from 2.5 dpf and impaired TD formation at 5 dpf (Fig. 1E-H), similar to *tie1* mutant larvae, although their phenotypes were milder than *tie1* mutants (see Fig. 1B,D). In contrast, neither *angpt2a* mutants nor *angpt2b* mutants exhibited these phenotypes (Fig. 1E-H). In addition, the phenotypes observed in *angpt1* mutants were not significantly exacerbated in triple-homozygous *angpt1*-*angpt2a*-*angpt2b* mutants, indicating that Angpt2a and Angpt2b do not compensate for the loss of Angpt1 in lymphatic development. Therefore, our results indicate that among Angiopoietins and Tie receptors, Angpt1 and Tie1 function in lymphatic development in the zebrafish trunk.

### Angpt1 acts as the Tie1 ligand in zebrafish

In zebrafish, *tie1* is expressed in all ECs in developing and mature blood vessels (Carlantoni et al., 2021; Lyons et al., 1998; Pham et al., 2001). We confirmed that *tie1* transcripts are present in the DA, PCV, and intersegmental vessels (ISVs) at 56 hpf by RNAscope analyses (Fig. S2A). On the other hand, *angpt1* is predominantly expressed in the hypochord (Fig. S2B), a single layer of cells immediately dorsal to the DA, as well as ventral mesenchyme surrounding the major trunk vessels (Lamont et al., 2010; Pham et al., 2001). Thus, *angpt1*-expressing cells are in close proximity to *tie1*-expressing ECs in the trunk; however, it has not been demonstrated whether Angpt1 directly binds to zTie1 as a ligand. To examine this, we performed an *in vitro* binding assay. We tested the association of immunoglobulin Fc-domain tagged soluble extracellular domain of zTie1 (zTie1-Fc) or zTie2 (zTie2-Fc) with Angpt1, Angpt2a or Angpt2b (Fig. 2A). zTie1-Fc specifically bound to Angpt1 but not to Angpt2a or Angpt2b (Fig. 2B), suggesting that Angpt1 serves as a ligand for zTie1. This specific binding of Angpt1 and zTie1 is consistent with the fact that *angpt1* mutants phenocopied *tie1* mutants (Fig. 1). We also detected interaction of zTie2-Fc with both Angpt1 and Angpt2a although the biological importance of their bindings is unclear (Fig. 2B).

**Figure 2.**
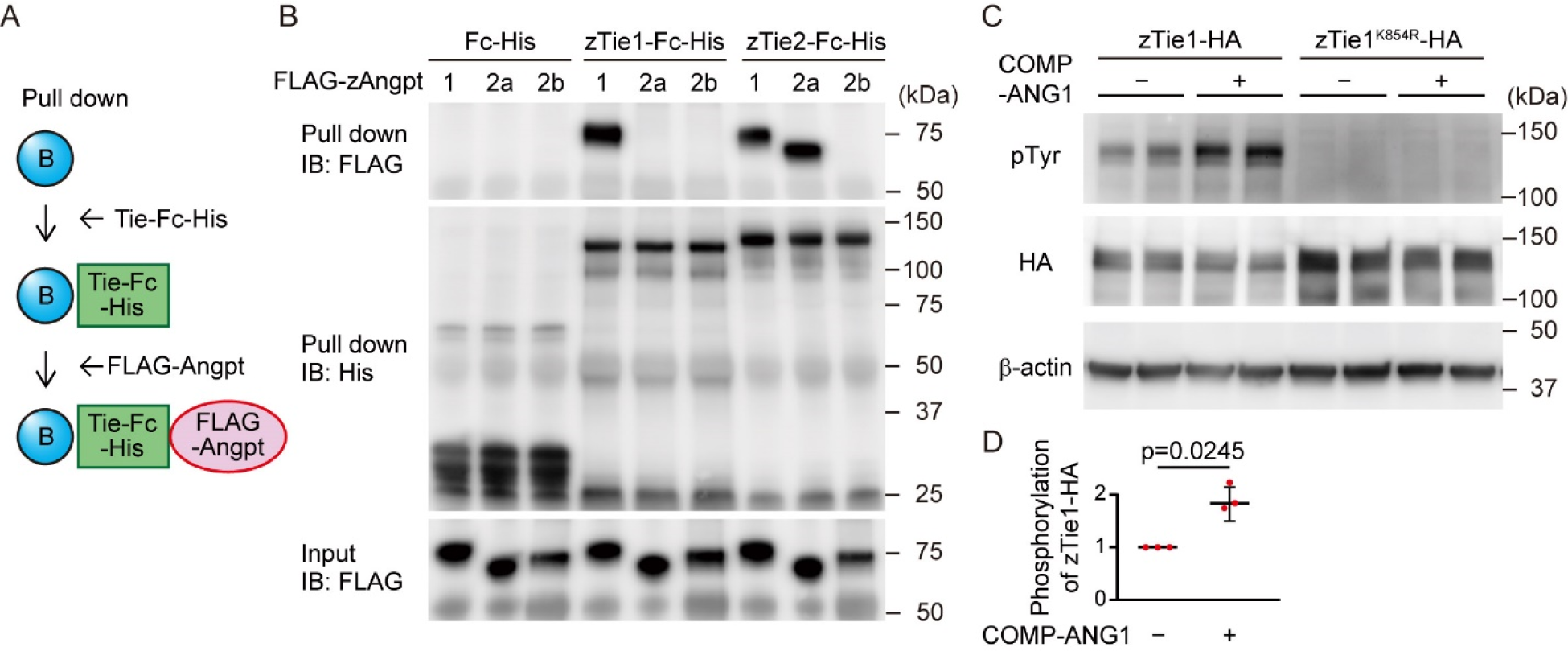
Zebrafish Tie1 (zTie1) acts as the receptor for zAngpt1. (**A**) Schematic illustration of the experiment for *in vitro* binding assay for (B). (**B**) *In vitro* binding of zebrafish Tie1 (zTie1)-Fc-His, zTie2-Fc-His or Fc-His. Binding in the presence of FLAG-Angpt1, FLAG-Angpt2a or FLAG-Angpt2b was examined as described in the ‘Materials and methods’ section. (**C**) 293T cells transfected with full length zTie1 (zTie1-HA) and a kinase-deficient mutant of zTie1 (zTie1^K854R^-HA) were starved and stimulated with vehicle (–) and COMP-Ang1 (+) for 20 minutes. Cell lysates were subjected to immunoblot analyses with anti-phosphotyrosine (pTyr) and anti-HA antibodies for analyzing phosphorylated Tie1 and total Tie1, respectively, and also with anti-β-actin antibody. (**D**) Relative phosphorylation of zTie1 observed in (C) was quantified. Data are mean ± s.d. from three independent experiments, each experiment includes duplicate determinations.

Next, we examined phosphorylation of zTie1 by Angiopoietin. Of note, upon stimulation with cartilage oligomeric matrix protein (COMP)-ANG1, a potent human Angiopoietin 1 variant (Cho et al., 2004), Tyr phosphorylation of full-length zTie1 but not kinase-dead zTie1 (zTie1^K854R^) was significantly enhanced (Fig. 2C,D). Therefore, zTie1 can function as a receptor tyrosine kinase that responds to Angiopoietin. Collectively, our results strongly suggest that Angpt1/Tie1 signaling functions in zebrafish.

### Angpt1/Tie1 signaling is important for sprouting from the vein at the onset of lymphangiogenesis

To determine which event in lymphangiogenesis is regulated by Angpt1/Tie1 signaling, we monitored trunk lymphatic development in live *tie1* mutants. In zebrafish, lymphangiogenesis starts with endothelial sprouting from the PCV, which is called secondary sprouting (Fig. S3A and Movie 1) (Hogan and Schulte-Merker, 2017; Isogai et al., 2003). About half of sprouts from the PCV connect to the primary ISVs to form veins (venous ISVs (vISVs)). The other half of the sprouts do not connect to the ISVs but migrate to the horizontal myoseptum to form parachordal lymphangioblasts (PLs), a pool of lymphatic precursors, which will form the entire trunk lymphatic vasculature including the TD (Fig. S3A and Movie 1). When we observed lymphatic development prior to TD formation, we found that PLs were mostly absent in *tie1* mutants at 54 hpf (Fig. 3A,B). Therefore, Tie1 is essential for proper development of lymphatic precursors in the trunk. We then examined the secondary sprouting from the vein. In *tie1* mutants, the number of secondary sprouts was markedly reduced in the trunk (Fig. 3A,C) as well as in the tail (Fig. S3B). Because the secondary sprouts contribute to both lymphatic and venous blood vessels, we examined whether Tie1 only regulates lymphatic development or both. In *tie1* mutants, both vISV and PL formation were blocked (Fig. 3A,B,D), indicating that Tie1 is essential for secondary sprouting that contributes to both lymphatic and venous vessel development. Moreover, arterial-venous (A-V) connections consisting of an arterial ISV (aISV) and a secondary sprout remained in the mutants at 54 hpf (Fig. S3C-E), indicating that Tie1 is also important for vessel remodeling to form vISVs. Similar to the *tie1* mutants, *angpt1* mutants also exhibited impaired PL formation and a reduced number of secondary sprouts and vISVs (Fig. S3F-I). These results demonstrate that Angpt1/Tie1 signaling regulates secondary sprouting from the large vein at the onset of lymphangiogenesis.

**Figure 3.**
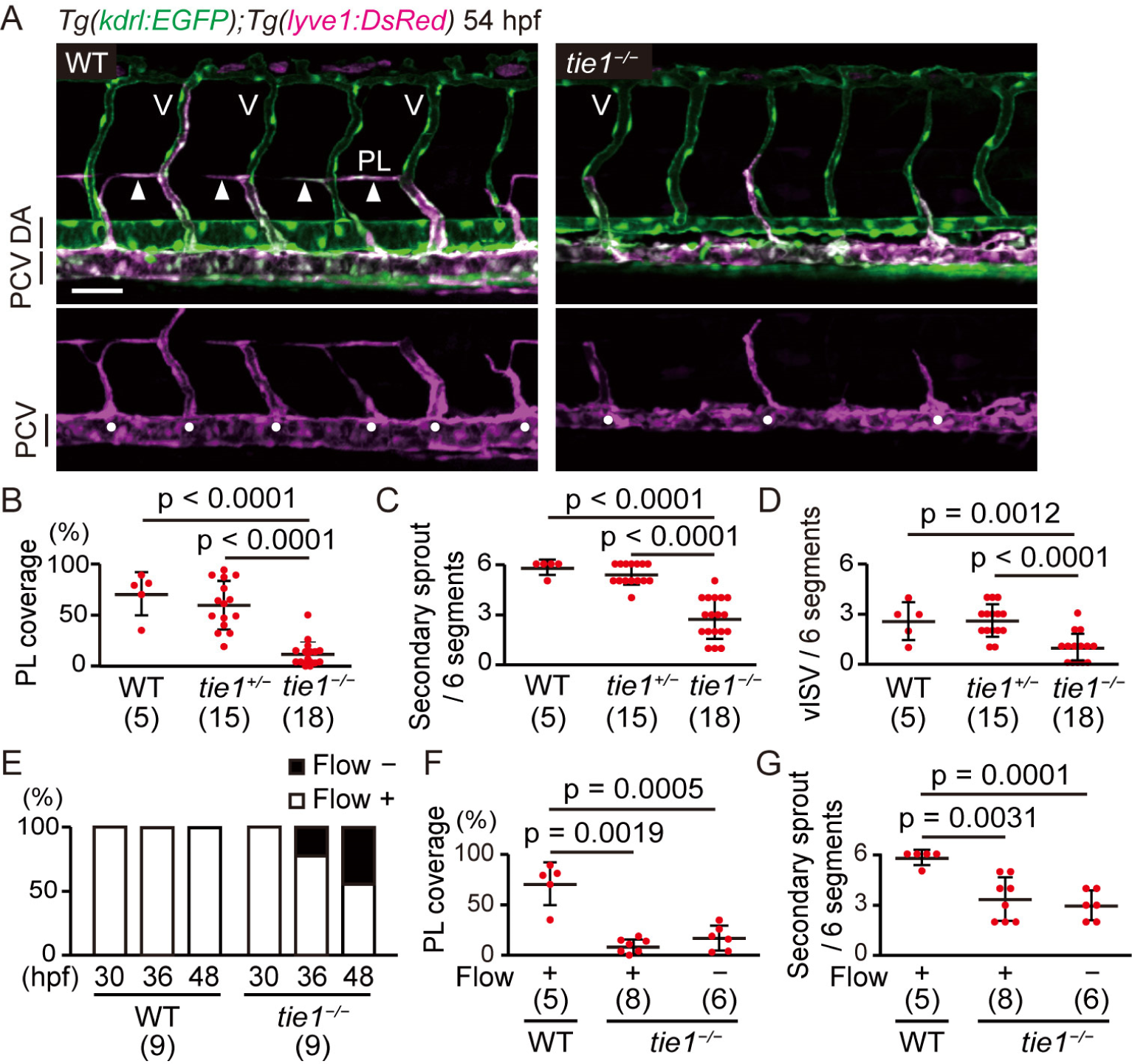
*tie1* mutants exhibit defects in secondary sprouting from the posterior cardinal vein (PCV). (**A**) Trunk of *Tg(kdrl:EGFP);Tg(lyve1:DsRed)* WT and *tie1^−/−^* embryos (54 hpf). Formation of secondary sprouts (solid circles), PLs (arrowheads), and venous ISVs (V) were inhibited in *tie1^−/−^*embryos. (**B**) Percentage of PL coverage at 54 hpf. Data are mean ± s.d. (WT, n = 5 embryos; *tie1^+/−^*, n = 15 embryos; *tie1^−/−^*, n = 18 embryos). (**C**) Number of secondary sprouts scored across 6 segments at 54 hpf. Data are mean ± s.d. (WT, n = 5 embryos; *tie1^+/−^*, n = 15 embryos; *tie1^−/−^*, n = 18 embryos). (**D**) Number of vISVs scored across 6 segments at 54 hpf. Data are mean ± s.d. (WT, n = 5 embryos; *tie1^+/−^*, n = 15 embryos; *tie1^−/−^*, n = 18 embryos). (**E**) Percentage of WT and *tie1^−/−^*embryos with (+) and without (−) blood flow at 30, 36, and 48 hpf. The blood flow in *tie1* mutants was gradually decreased during and after secondary sprouting (WT, n = 9 embryos; *tie1^−/−^*, n = 9 embryos). (**F**) Percentage of PL coverage in the trunk of WT and *tie1^−/−^* embryos with (+) and without (−) blood flow at 54 hpf. Data are mean ± s.d. (WT, n = 5 embryos; blood flow (+) *tie1^−/−^*, n = 8 embryos; blood flow (−) *tie1^−/−^*, n = 6 embryos). (**G**) Number of secondary sprouts scored across 6 segments at 54 hpf. Data are mean ± s.d. (WT, n = 5 embryos; blood flow (+) *tie1^−/−^*, n = 8 embryos; blood flow (−) *tie1^−/−^*, n = 6 embryos). Scale bar: 50 μm. PL, parachordal lymphangioblast.

*tie1* mutants gradually lost blood flow from 36 hpf (Fig. 3E), most likely due to cardiac defects (Carlantoni et al., 2021). Because blood flow is involved in lymphatic vessel formation (Coffindaffer-Wilson et al., 2011), we examined whether the phenotypes in *tie1* mutants might be caused by the loss of blood flow. With or without blood flow, *tie1* mutants similarly exhibited impaired PL formation and a reduced number of secondary sprouts (Fig. 3F,G). Although secondary sprouting was partially blocked by stopping blood flow with Verapamil treatment, the effects were much milder than those in *tie1* mutants (Fig. S3J). Therefore, impaired blood flow is not the main cause of the phenotypes in *tie1* mutants. In addition, blood flow was mostly unaffected in *angpt1* mutants that exhibit similar phenotypes to *tie1* mutants (Fig. S3K). Therefore, Angpt1/Tie1 signaling regulates lymphangiogenesis independently of flow regulation.

### Tie1 signaling is important for lymphatic specification, migration, and proliferation

Next, we investigated cell behaviors controlled by Tie1 signaling during secondary sprouting. In *tie1* mutants with reduced numbers of secondary sprouts (Fig. 3C), ECs extend their protrusions, but their nuclei remained mostly within the parental vein and rarely crossed the level of the DA (Fig. 4A,B and Movie 2). Therefore, Tie1 is important for the migration of EC nuclei from the PCV. In addition, even when their nuclei migrate dorsally, their migration distance and speed were significantly lower than those of wild-type siblings (Fig. 4C,D). Therefore, Tie1 is important for the migration of budding ECs from the PCV.

**Figure 4.**
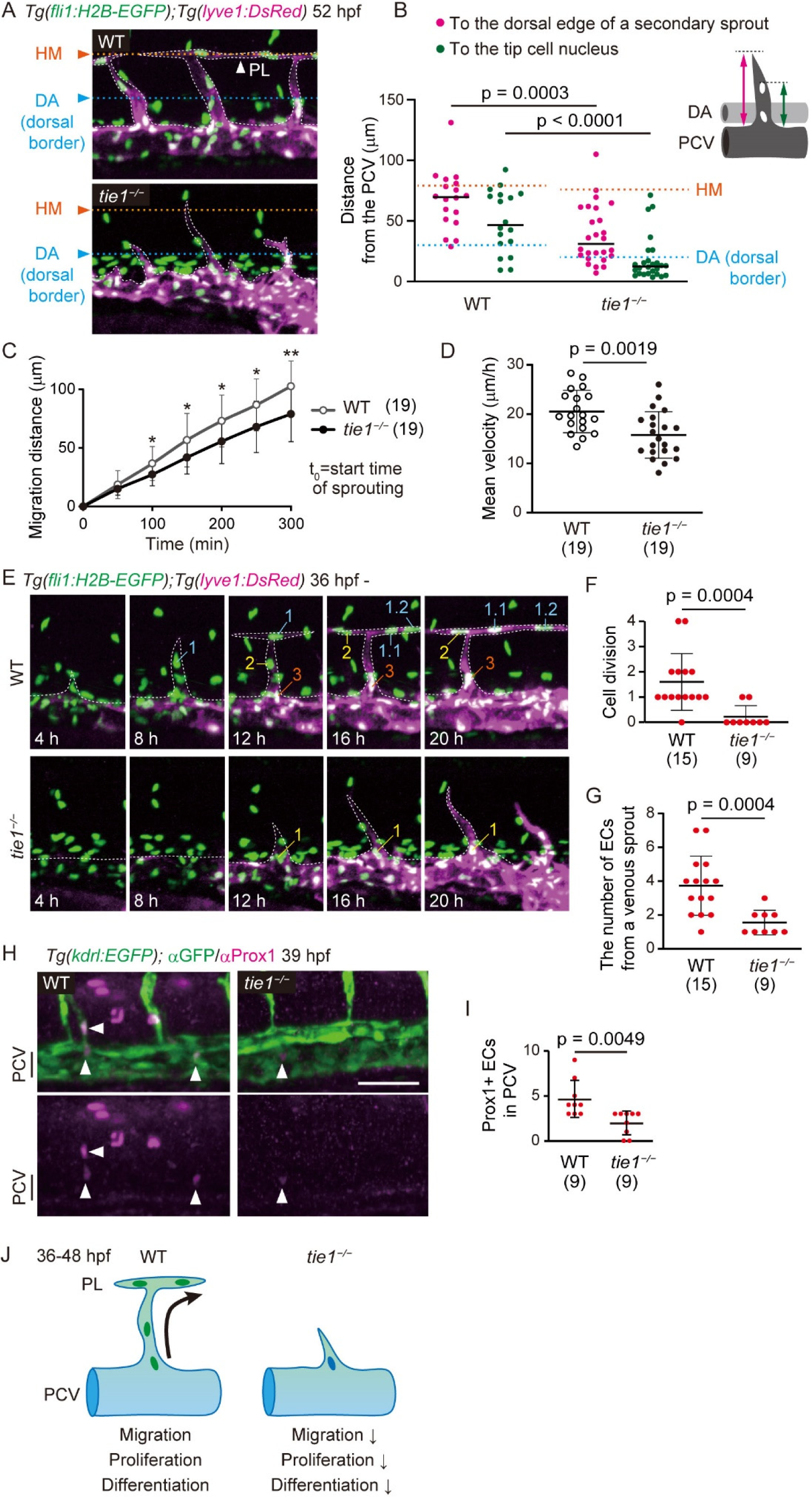
zTie1 regulates migration, proliferation, and differentiation of sprouting ECs from the PCV. (**A**) Trunk of *Tg(fli1:H2B-EGFP);Tg(lyve1:DsRed)* WT and *tie1^−/−^* embryos (54 hpf). Orange, cyan, and white dot lines indicate horizontal myoseptums (HM), dorsal borders of the dorsal aorta (DA), and outlines of *lyve1*:DsRed-positive venous ECs, respectively. (**B**) Distance from the dorsal edge of the PCV to the dorsal edge of a secondary sprout (magenta dots) and to the tip cell nucleus (green dots) in WT and *tie1^−/−^* embryos at 52 hpf. Data are mean ± s.d. (WT, n = 18 sprouts in 4 embryos; *tie1^−/−^*, n = 26 sprouts in 7 embryos). (**c**) Total distance traveled by the tip cell nucleus at each time point from the initiation of venous sprouting. Start time of sprouting (t = 0) was 40.0 ± 3.7 hpf in WT and 42.7 ± 4.5 hpf in *tie1* mutant. Data are mean ± s.d. (WT, n = 19 sprouts in 3 embryos; *tie1^−/−^*, n = 19 sprouts in 4 embryos). *P < 0.05; **P < 0.01. (**D**) Mean velocity of the tip cell nucleus during the first 300 minutes of venous sprouting. Data are mean ± s.d. (WT, n = 19 sprouts in 3 embryos; *tie1^−/−^*, n = 19 sprouts in 4 embryos). (**E**) Time-sequential images of the trunk of *Tg(fli1:H2B-EGFP);Tg(lyve1:DsRed)* WT and *tie1^−/−^* embryos from 36 hpf (when secondary sprouting starts in WT). White dot lines outline *lyve1*:DsRed-positive ECs. (**F**) Number of cell division in a venous sprout from 36 to 56 hpf. Data are mean ± s.d. (WT, n = 15 sprouts in 4 embryos; *tie1^−/−^*, n = 9 sprouts in 7 embryos). (**G**) Number of ECs from a venous sprout at 56 hpf. Data are mean ± s.d. (WT, n = 15 sprouts in 4 embryos; *tie1^−/−^*, n = 9 sprouts in 7 embryos). (**H**) Whole-mount immunofluorescence staining for Prox1 (magenta) and GFP (green) in WT and *tie1^−/−^ Tg(kdrl:EGFP)* embryos at 39 hpf. Arrowheads indicate Prox1-positive ECs. (**I**) Number of Prox1-positive ECs in the PCV across 6 segments at 39 hpf. Data are mean ± s.d. (WT, n = 9 embryos; *tie1^−/−^*, n = 9 embryos). (**J**) During the process of lymphangiogenesis, ECs budding from the PCV migrate, proliferate, and undergo lymphatic differentiation. In contrast, all of these cell behaviors are inhibited in *tie1* mutants. Scale bar: 50 μm. HM, horizontal myoseptum.

Secondary sprouting ECs proliferate to form the PL and the vISVs (Fig. 4E). In contrast, in *tie1* mutants, such proliferation events were markedly reduced (Fig. 4E,F). Similarly, the number of ECs constituting each venous sprout was markedly reduced (Fig. 4G). Apoptosis was not enhanced in these mutants (Fig. S4). Therefore, Tie1 is important for the proliferation of budding venous ECs but dispensable for their survival.

Next, we examined the role of Tie1 in lymphatic differentiation. Prox1 starts to be expressed in lymphatic precursors of the PCV in a manner dependent on Erk activation induced by Vegfc/Vegfr3 signaling (Koltowska et al., 2015; Shin et al., 2016). Prox1^+^ ECs were found in the PCV and the secondary sprouts in the wild-type situation (Fig. 4H). In contrast, the number of Prox1-positive ECs was significantly reduced in *tie1* mutants (Fig. 4I). Consistent with this, phosphorylation of ERK was reduced in the secondary sprouts of the *tie1* mutants (see Figure 7A). Therefore, Tie1 regulates the induction of Prox1 presumably by affecting ERK phosphorylation. Taken together, these results demonstrate that Tie1 signaling regulates multiple EC behaviors in lymphangiogenesis: EC migration, proliferation, and lymphatic differentiation (Fig. 4J).

### Tie1 signaling regulates gene expression involved in lymphatic development

To investigate what kind of gene expression is regulated downstream of Tie signaling, we performed RNA sequencing (RNA-seq) analyses of ECs from wild-type and *tie1* mutant embryos during secondary sprouting. To this end, *kdrl*:EGFP/*lyve1*:DsRed double-positive venous and early lymphatic ECs were isolated from the trunk and tail of embryos at 47 hpf by fluorescence-activated cell sorting (FACS) and were subjected to RNA-seq analyses (Fig. 5A). Volcano plot analysis showed that the number of down-regulated genes in the *tie1* mutants was much greater than the number of up-regulated genes (Fig. 5B). We identified 1,444 differentially expressed genes (DEGs) with q-value < 0.05 between homozygous *tie1* mutant embryos and their wild-type and heterozygous siblings (Fig. 5C). Among the DEGs, 1416 genes were downregulated in *tie1* mutants (Table S1). Genes down-regulated in the *tie1* mutants included those related to blood and lymphatic vascular development, such as *aplnra*, *lmo2*, and *sox18* (Fig.5 D-F). Downregulated genes are also enriched for those that function in cell migration and cell cycle progression (Fig. 5E,F), consistent with impaired migration and proliferation in *tie1* mutants (Fig. 4). These results suggest that Tie1 signaling controls a series of gene expression events involved in lymphatic development.

**Figure 5.**
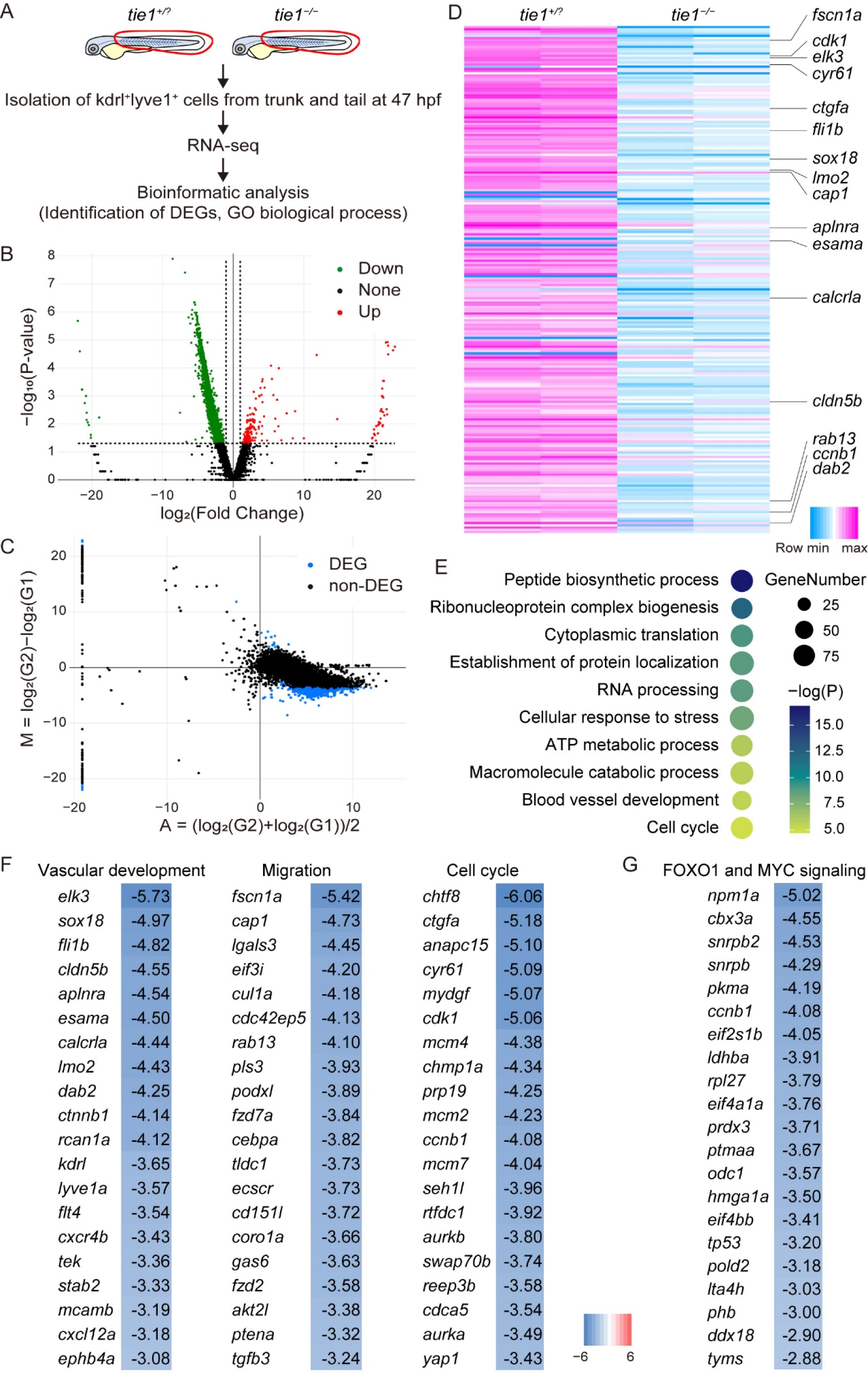
Tie1 signaling regulates gene expression involved in lymphatic development. (**A**) Schematic of the workflow. RNA sequencing (RNA-seq) was performed in *tie1^−/−^* embryos and their *tie1*^+/?^ siblings (n = 2 RNA samples each). (**B**) The volcano plots of the *tie1^−/−^* and *tie1^+/?^* gene set. The red, green, and black dots represent up-regulated (in *tie1^−/−^* embryos), down-regulated and no significance genes, respectively, with fold change > 1 and p-value < 0.05. (**C**) The MA plots of 1,444 DEGs (blue dots) with q-value < 0.05 in the *tie1^−/−^* and *tie1^+/?^* gene set. (**D**) Heatmap of the most significant DEGs using log2(TPM) values. (**E**) Representative GO biological processes of the DEGs. (**F**) Heatmap of the DEGs associated with the GO terms “Vascular development”, “Migration”, and “Cell cycle.” Genes are ranked by log2 fold change *tie1^−/−^*/*tie1^+/?^*. (**G**) Heatmap of the DEGs associated with the FOXO1 and MYC signaling. Genes are ranked by log2 fold change *tie1^−/−^*/*tie1^+/?^*.

Of note, genes down-regulated in *tie1* mutants included those associated with forkhead box O (FOXO) transcription factor FOXO1 (Fig. 5G), known as a gatekeeper of endothelial quiescence (Wilhelm et al., 2016). Endothelial FOXO1 suppresses signaling by MYC. MYC signature genes repressed by nuclear FOXO1 (Wilhelm et al., 2016) were similarly downregulated in the *tie1* mutants (Figure 5G). Therefore, FOXO1 might be used as a readout of Angpt1/Tie1 signaling.

### Tie1 inhibits nuclear import of Foxo1a through the PI3K pathway in secondary sprouting ECs

In ECs, FOXO1 is excluded from the nucleus by AKT-mediated phosphorylation downstream of phosphatidylinositol-3-OH kinase (PI3K) signaling (Eijkelenboom and Burgering, 2013). To test whether FOXO1 is regulated downstream of Tie1, we examined the localization of FOXO1 by generating a transgenic fish (Tg) line, *Tg(fli1:EGFP-Foxo1a)*, which expresses EGFP-tagged zebrafish Foxo1a specifically in ECs. We simultaneously marked EC nuclei, by crossing with *Tg(fli1:H2B-mC)*, in which EC nuclei were labeled by histone H2B-mC (Yokota et al., 2015). In the PCV and caudal vein plexus (CVP), endothelial EGFP-Foxo1a was predominantly detected in the cytoplasm in wild-type at 31 hpf before secondary sprouting (Fig. 6A). In contrast, in *tie1* mutants, nuclear localization of EGFP-Foxo1a was clearly enhanced, especially in the CVP (Fig. 6A), indicating that Tie1 is important for nuclear exclusion of EGFP-Fox1a, at least in some venous ECs before secondary sprouting. During secondary sprouting, EGFP-Foxo1a was excluded from the nucleus in most of sprouting ECs of wild-type and heterozygous *tie1^+/−^* embryos (Fig. 6B,C). Even when EGFP-Foxo1a was first localized in the nucleus prior to sprouting, it was exported from the nucleus before or during the sprouting (Fig. 6D). However, EGFP-Foxo1a accumulated in the nuclei of sprouting ECs in *tie1^−/−^* embryos (Fig. 6B,C), where their nuclear migration was significantly blocked (Fig. 4A,B). These results indicate that Tie1 signaling inhibits nuclear import of EGFP-Foxo1a in the sprouting venous ECs. To explore whether Tie1-dependent nuclear exclusion of EGFP-Foxo1a is mediated by the PI3K/Akt pathway, we examined the effect of PI3K signaling inhibition. Treatment with LY294002, a PI3K inhibitor, altered the localization of EGFP-Foxo1a in secondary sprouting ECs from cytoplasmic to nuclear (Fig. 6E). Therefore, nuclear exclusion of EGFP-Foxo1a during secondary sprouting is mediated by PI3K. Furthermore, after EGFP-Foxo1a entered the nuclei, these budding ECs retracted into the PCV (Fig. 6E; in 7/8 ECs), highlighting the importance of the PI3K pathway in secondary sprouting. Collectively, our results indicate that Tie1 signaling is activated in secondary sprouting ECs and inhibits nuclear import of FOXO1 through the PI3K pathway.

**Figure 6.**
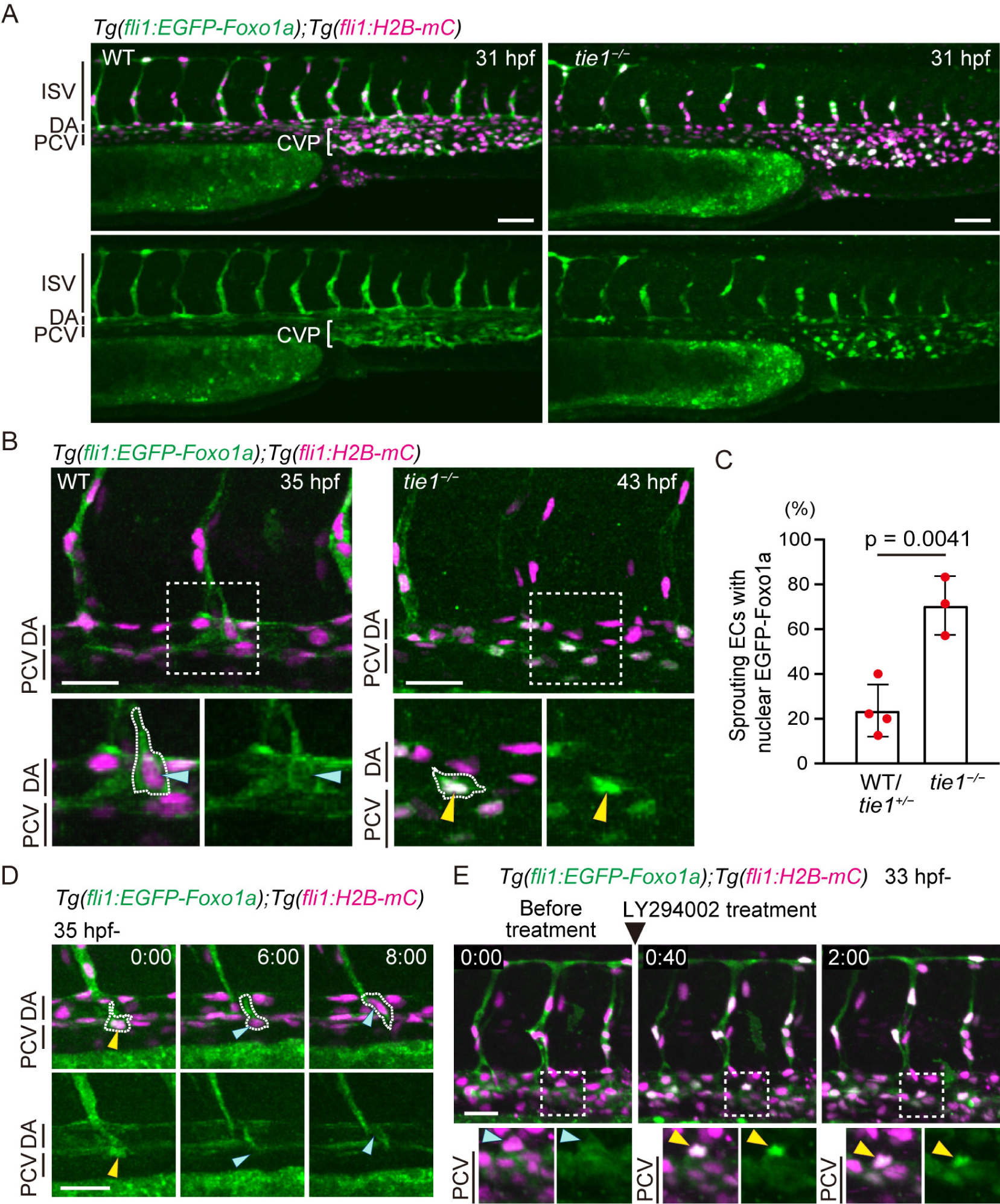
Tie1 inhibits nuclear import of Foxo1a during secondary sprouting. (**A**) Trunk and tail of *Tg(fli1:EGFP-Foxo1a);Tg(fli1:H2B-mCherry)* WT and *tie1^−/−^* embryos (31 hpf). (**B**) Trunk of *Tg(fli1:EGFP-Foxo1a);Tg(fli1:H2B-mCherry)* WT (35 hpf) and *tie1^−/−^* embryos (43 hpf). The boxed areas are enlarged in the bottom. White dot lines outline ECs sprouting from the PCV. Blue and yellow arrowheads indicate EGFP-Foxo1a in the cytoplasm and nucleus, respectively. (**C**) Quantitative analysis of the data shown in (B). The graph shows the percentage of secondary sprouting ECs with nuclear EGFP-Foxo1a among total sprouting ECs from the PCV and CVP. Each dot represents an individual embryo. Data are mean ± s.d. (WT/*tie1*^+/−^, n = 4 embryos; *tie1^−/−^*, n = 3 embryos). 6-10 sprouting ECs were measured in each embryo. (**D**) Time-sequential images of the trunk of a *Tg(fli1:EGFP-Foxo1a);Tg(fli1:H2B-mCherry)* WT embryo from 35 hpf. Elapsed time (h:min). White dot lines outline an EC sprouting from the PCV. Before nuclear migration, EGFP-Foxo1a was exported from the nuclei (yellow arrowheads) to the cytoplasm (blue arrowheads). (**E**) Time-sequential images of the trunk of a *Tg(fli1:EGFP-Foxo1a);Tg(fli1:H2B-mCherry)* embryo (from 33 hpf) treated with 30 μM LY294002 just after the z-stack imaging at 0:00. Elapsed time (h:min). The boxed areas are enlarged in the bottom. Treatment with LY294002 altered the localization of EGFP-Foxo1a in a secondary sprouting EC from cytoplasmic (blue arrowheads) to nuclear (yellow arrowheads), leading to retraction of the sprouts. Scale bars: 50 μm (A), 30 μm (B,D,E). CVP, caudal vein plexus.

### Tie1 signaling regulates lymphangiogenesis through the modulation of Vegfc/Vegfr3 signaling

We finally examined the relationship between Angpt1/Tie1 signaling and Vegfc/Vegfr3 signaling in lymphatic development. Vegfc/Flt4 signaling induces phosphorylation of ERK, thereby leading to lymphatic differentiation and secondary sprouting in the trunk of zebrafish (Koltowska et al., 2015; Shin et al., 2016). In *tie1* mutants, phosphorylation of ERK was significantly reduced in secondary sprouting ECs (Fig. 7A,B). This result suggests that Vegfc/Flt4/Erk signaling is attenuated in *tie1* mutants. To determine whether the *tie1* mutant phenotypes could be explained by reduced Vegfc/Vegfr3 signaling, we examined whether enhancing Vegfc/Vegfr3 signaling could rescue the *tie1* mutant phenotypes. When *vegfc* was overexpressed in the arterial ECs of *tie1* mutant (using *Tg(flt1:Gal4FF)*;*Tg(10xUAS:Vegfc)* double transgenic embryos) (Koltowska et al., 2015), the phenotypes of decreased secondary sprouts, vISV formation, and PL coverage in *tie1* mutants were partially rescued (Fig. 7C-F). These results indicate that the impaired lymphatic development in *tie1* mutant can be explained, at least in part, by reduced responsiveness of Vegfc/Vegfr3 signaling.

**Figure 7.**
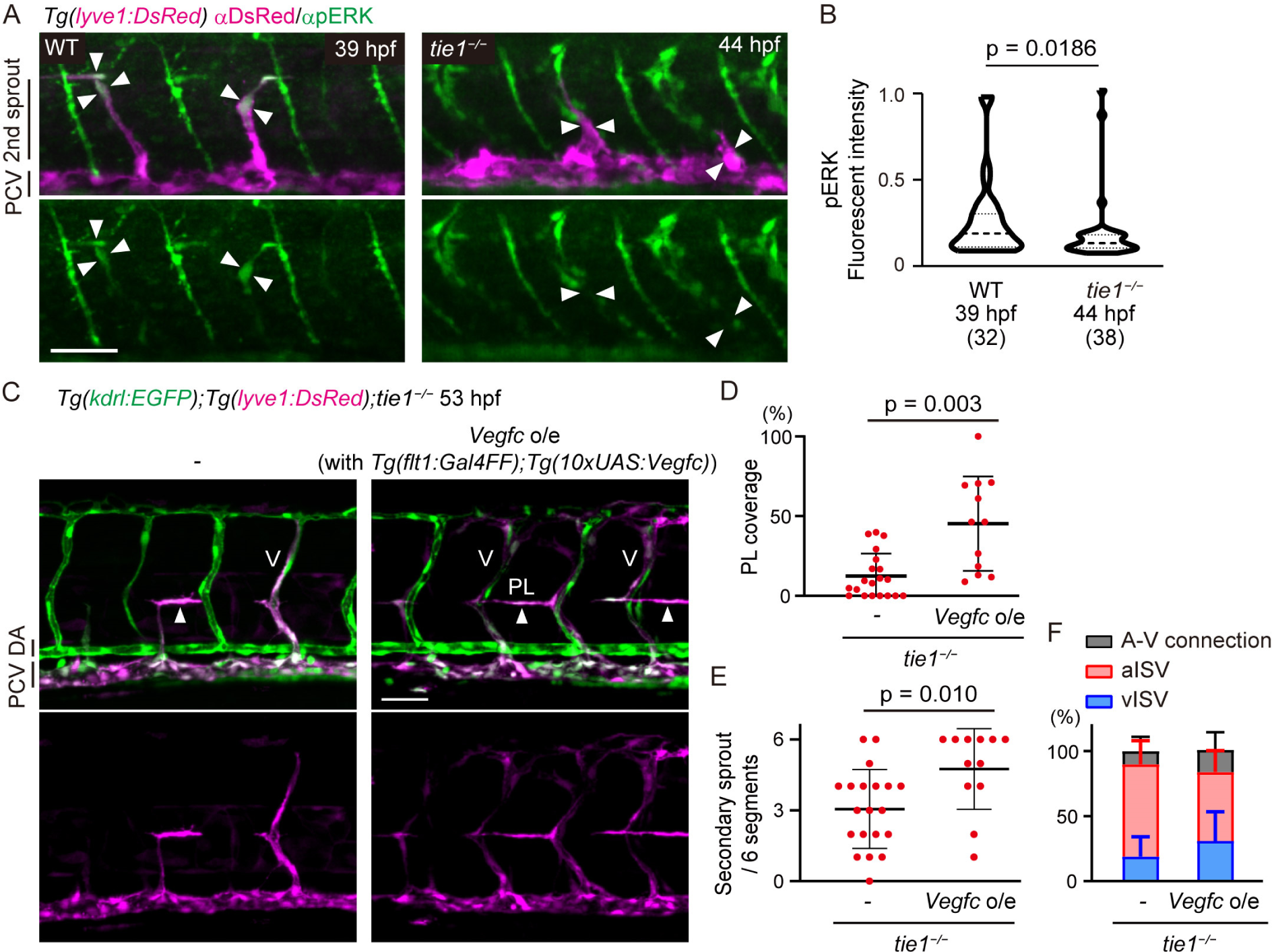
Tie1 regulates lymphatic development at least in part through the modulation of Vegfc/Vegfr3 signaling. (**A**) Whole-mount immunofluorescence staining for phospho ERK (green) and DsRed (magenta) in WT and *tie1^−/−^ Tg(lyve1:DsRed)* embryos. Arrowheads point to EC nuclei of secondary sprouts assessed by the DAPI staining. Secondary sprouts migrate towards the HM around 39 hpf in WT and 44 hpf in *tie1* mutant. (**B**) Quantitative analyses of the data shown in (A). Graph shows fluorescent intensity of pERK staining in ECs budding from the PCV of WT (39 hpf) and *tie1^−/−^* (44 hpf) embryos. Data are mean ± s.d. (WT, n = 32 ECs in 5 embryos; *tie1^−/−^*, n = 38 ECs in 7 embryos). (**C**) Trunk of *Tg(kdrl:EGFP);Tg(lyve1:DsRed) tie1^−/−^* embryos with or without *Tg(flt1:Gal4FF);Tg(10×UAS:vegfc)* (53 hpf). Formation of PLs (arrowheads) and venous ISVs (Vs) are partially recovered by overexpression of Vegfc in *tie1* mutant. (**D**) Percentage of PL coverage at 54 hpf. Data are mean ± s.d. (*tie1^−/−^*, n = 20 embryos; *tie1^−/−^* with Vegfc o/e, n = 12 embryos). (**E**) Number of secondary sprouts scored across 6 segments at 54 hpf. Data are mean ± s.d. (*tie1^−/−^*, n = 20 embryos; *tie1^−/−^* with Vegfc o/e, n = 12 embryos). (**F**) Number of A-V connection across 6 segments at 54 hpf. Data are mean ± s.d. (*tie1^−/−^*, n = 20 embryos; *tie1^−/−^* with Vegfc o/e, n = 12 embryos). Scale bars: 50 μm.

## DISCUSSION

In the present study, we performed comprehensive analyses of *angiopoietin* and *tie* mutants in zebrafish and thereby found that Angpt1 and Tie1 are important for trunk lymphatic development. In contrast, we could not find any roles of Angpt2a, Angpt2b, and Tie2, and mutants in either gene did not show any additional phenotypes. We also demonstrate the binding between Angpt1 and zTie1 by *in vitro* binding assays. Thus, our results demonstrate that Angpt1 can serve as a ligand for zTie1. In addition, we show that zTie1 functions as a RTK that undergoes auto-phosphorylation upon Angiopoietin stimulation. Therefore, our results show that Angpt1/Tie1 signaling functions in zebrafish lymphatic development.

*angpt1* mutants and *tie1* mutants exhibit similar phenotypes including reduced secondary sprouting, decreased vISV and PL formation, as well as cardiac edema. However, all of those phenotypes are milder in *angpt1* mutants than in *tie1* mutants. Considering that Angpt2a and Angpt2b do not compensate for the loss of *angpt1*, it is suggested that molecules other than Angiopoietin might regulate Tie1 cooperatively or independently of Angpt1. It has been recently shown (Hußmann et al., 2023) that Tie1 has physical and genetic interaction with Svep1 (also known as Polydom), an extracellular matrix indispensable for lymphatic development in mice and zebrafish (Hußmann et al., 2023; Karpanen et al., 2017; Morooka et al., 2017; Sato-Nishiuchi et al., 2023). Similar to *tie1* mutants, *svep1* mutant fish exhibit defects in the trunk lymphatic development (Karpanen et al., 2017; Morooka et al., 2017). In addition, a recent report shows that mammalian SVEP1 can promote LEC migration via TIE1, but without apparent TIE1 phosphorylation (Sato-Nishiuchi et al., 2023). Therefore, Svep1 is a promising candidate for Tie1 activators other than Angpt1. Since Ang1 binds to Svep1 in mice (Morooka et al., 2017), it is possible that Angpt1 and Svep1 coordinately regulate Tie1 signaling to control proper lymphatic development.

We here show that Angpt1/Tie1 signaling and Vegfc/Vegfr3 signaling cooperatively regulate the development of trunk lymphatic vessels, including the PL, TD, and the dorsal longitudinal lymphatic vessel (DLLV). Tie1 regulates trunk lymphatic development at least in part through the modulation of Vegfc signaling. On the other hand, in facial lymphatic development, Tie1 and Vegfc signaling act separately in a different context of lymphatic vessel formation (Hußmann et al., 2023). Vegfc regulates lymphatic branchial arches, lateral and medial facial lymphatics, and otolithic lymphatic vessels independently of Tie1, whereas Tie1, together with Svep1, regulates the facial collecting lymphatic vessel (FCLV) development (Hußmann et al., 2023). These results indicate that regulation of lymphatic development varies in different lymphatic vessels, and additional efforts will have to be made to understand how Vegf signaling and Tie1 signaling, sometimes in concert and sometimes independently, regulate lymphangiogenesis. To clarify this, it will be important to elucidate the downstream molecular mechanisms of each signaling pathway.

Tie1 plays a crucial role in PI3K-mediated nuclear exclusion of Foxo1a in secondary sprouting ECs in zebrafish. Wilhelm et al, report that murine Foxo1 acts as a gatekeeper of endothelial quiescence, which decelerates metabolic activity through c-Myc inhibition (Wilhelm et al., 2016). Consistently, our RNA-seq results identify that Tie1 signaling positively regulates the expression of Myc target genes and metabolic-related genes. Therefore, the Tie1/PI3K signaling pathway may regulate lymphatic development through changes in metabolic state of ECs. It would be interesting to investigate the importance of metabolic regulation in lymphangiogenesis that might be regulated by Angpt1/Tie1 signaling.

In zebrafish that lack functional Tie2, Tie1 plays an essential role in trunk lymphangiogenesis. Similarly in mammals, TIE1 appears to have a unique role in lymphatic vascular development (D’Amico et al., 2010; Donnan et al., 2021; Qu et al., 2010; Qu et al., 2015; Shen et al., 2014); however, how mammalian TIE1 regulates lymphangiogenesis together with TIE2 or independently of TIE2 is not fully understood. It will be an interesting challenge to investigate how Tie1 signaling has been altered or conserved during the evolution from fish to mammals.

## MATERIALS AND METHODS

### Zebrafish husbandry and strains

Zebrafish (*Danio rerio*) were maintained and bred under standard conditions (Alestrom et al., 2020). The experiments using zebrafish were approved by the animal committee of the National Cerebral and Cardiovascular Center (No.22054) and performed according to our institutional regulation. The following transgenic and mutant zebrafish lines were used for this study: *Tg(fli1:EGFP)^y1^* (Lawson and Weinstein, 2002), *Tg(kdrl:EGFP)^s843^* (Jin et al., 2005), *Tg(−5.2lyve1b:DsRed)^nz101^* (Okuda et al., 2012), *Tg(fli1:H2B-GFP)^ncv69^* (Ando et al., 2019), *Tg(fli1:H2B-mCherry)^ncv31^* (Yokota et al., 2015), *Tg(flt1:Gal4FF, myl7:Nls-mCherry)^ncv555^* (abbreviated as *Tg(flt1:Gal4FF)*) and *tie2^hu1667^* (Gjini et al., 2011). *Tg(flt1:Gal4FF)^ncv555^* and *Tg(fli1:EGFP-Foxo1a)^ncv557^* lines were generated by injecting the pTol flt1:Gal4FF plasmid and pTol fli1:EGFP-Foxo1a plasmid with Tol2 transposase mRNA into one cell stage embryos, respectively, as described previously (Nakajima et al., 2023). *Tg(10×UAS:Vegfc)^ncv556^* line was established using the same plasmid used to generate *Tg(10×UAS:Vegfc)^uq2bh^* (Koltowska et al., 2015). Throughout the text, all Tg lines used in this study are simply described without their line numbers. For example, *Tg(fli1:H2B-GFP)^ncv69^* is abbreviated to *Tg(fli1:H2B-GFP)*.

The knockout alleles ncv110 for *angpt1*, ncv129 for *angpt2a*, ncv130 for *tie1*, and ncv131 for *angpt2b* genes were generated by TALEN techniques as described below.

### Generation of knockout zebrafish by TALEN

TALENs targeting *tie1*, *angpt1*, *angpt2a*, and *angpt2b* were designed using TAL Effector Nucleotide Targeter 2.0 software (Doyle et al., 2012) and constructed using the Golden Gate assembly method (Cermak et al., 2011). The TALENs were cloned into an RCIscript-GoldyTALEN vector (Addgene). TALEN mRNAs were *in vitro* transcribed from SacI-linearized expression plasmids with T3 RNA polymerase using a mMessage mMachine mRNA kit (Thermo Fisher). Embryos, injected with 30-60 pg of the TALEN mRNAs at one-cell stage, were raised to adulthood and crossed with wild-type AB to identify germline-mutated founders.

Screening for founders was conducted by genomic PCR and subsequent sequencing using the following primer sets: 5′-TCTTCCAGATGCTGTCATGG-3′ and 5′-ATCCCACTGTGGTCAAAACC-3′ for *tie1*; 5′-TGTGAGTTTTCCGTCCCATC-3′ and 5′-ATAACCGTGTAATCATCCAG-3′ for *angpt1*; 5′-GAGCAAATATGTTGAGATCATGGA-3′ and 5′-CTTTTGCAGCCACTGTGTGT-3′ for *angpt2a*; 5′-CTGCCGTGTCTGTGCTACCT-3′ and 5′-AAGCACCATGACTATTTTCCTTG-3′ for *angpt2b*. For genotyping mutants, PCR analyses of genomic DNAs were routinely performed using the same primer set and these amplified PCR products were analyzed using an MCE-202 MultiNA microchip electrophoresis system (Shimadzu) with the DNA-500 reagent kit (Shimadzu).

### Image acquisition and image processing

For imaging, the pigmentation of embryos was suppressed by the addition of 1-phenyl-2-thiourea (PTU) (Sigma-Aldrich) in E3 media. Embryos were dechorionated and mounted in 1% low-melting agarose poured on a 35-mm-diameter glass-base dish (Asahi Techno Glass), and then anesthetized in 0.02% tricaine in E3 medium. Confocal images were taken with a FluoView FV1000, FV1200 and FV3000 confocal upright microscope system (Olympus) equipped with water-immersion XLUMPlan FL N 20x/1.00 NA objective lenses (Olympus) and a multi-alkali or GaAsP photomultiplier tube regulated with FluoViewASW software (Olympus). The 405 nm, 473 nm, and 559 nm laser lines were used. Images were acquired sequentially to avoid cross-detection of the fluorescent signals. Image files were processed and analyzed with FV10-ASW4.2 viewer (Olympus), ImageJ (Schindelin et al., 2012), and IMARIS 9.2.1, 9.5.1, or 9.9.1 software (Oxford Instruments)..

### Immunohistochemistry

Whole-mount immunostaining of zebrafish embryo was performed in accordance with the methods of Le Guen et al. (2014) with some modifications (Le Guen et al., 2014). Briefly, embryos were fixed with 4% paraformaldehyde (PFA) in PBS for 1 hour at room temperature, and washed three times in ice-cold MeOH, and then incubated 1 hour in 3% H_2_O_2_ in MeOH on ice. Embryos were washed three times in ice-cold MeOH and kept in MeOH at −20°C for 1 week. The fixed tissues were washed three times in PBS containing 0.2% Triton X-100 (PBS-T) for 10 min at room temperature, blocked in PBS-T containing 1% bovine serum albumin (BSA) for 1 hour at 4°C, and incubated with primary antibodies for 2 days at 4°C. After six washes in PBS-T for 30 min at 4°C, the samples were incubated with Alexa Fluor conjugated secondary antibodies overnight at 4°C. After another six washes in PBS-T for 30 min at 4°C, the embryos were mounted in 1% low-melting agarose poured on a 35-mm-diameter glass-base dish (Asahi Techno Glass), and were analyzed with a Fluoview FV3000 (Olympus) confocal microscope. Primary antibodies used were rabbit anti-Prox1 (1:500, Angiobio), rabbit anti-Phospho-p44/42 MAPK (Erk1/2) (1:500, Cell signaling), and mouse anti-RFP (1:500, MBL). Secondary antibodies used were anti-rabbit Alexa Fluor 488 and anti-mouse Alexa Fluor 546 (1:1000, Invitrogen). TUNEL assay was performed using In situ cell death detection kit, fluorescein (Roche) according to the manufacturer’s instructions. The embryos were counterstained with DAPI during the fifth wash following secondary antibody staining.

### RNAscope

Embryos were fixed by 10% NBF for overnight at room temperature. After fixation, the solution was changed to 50% Methanol/PBS-Tw (PBS with 0.1% Tween 20) for 10 min, then changed to 100% Methanol at room temperature, and then stored in 100% Methanol at −20°C overnight. After rehydration, embryos were washed three-times for 10 min in PBS-Tw containing 1% BSA. Embryos were treated with protease plus for 40 min at 40°C, and subsequently washed three times for 10 min in PBS-Tw. For probe hybridization, embryos were treated in the *tie1*-C1 and *angpt1*-C3 probes (ACDbio) for overnight at 40°C. Embryos were washed and subjected to hybridization following the workflow in ACDbio. Embryos were stored in PBS-Tw at 4°C prior to confocal imaging.

### Plasmids

cDNA fragments encoding full-length zebrafish *tie1* and *tie2* were amplified by PCR using cDNA library from 72 hpf zebrafish embryo as a template, then inserted into pCS2+ vector (Clontech) fused with C-terminal HA tag, namely pCS2+-zTie1-HA and pCS2+-zTie2-HA plasmids. pCS2+-zTie1KD-HA vector encoding a kinase-deficient mutant of *tie1* (K854R) was generated using inverse PCR method with KOD-plus DNA Polymerase (TOYOBO).

To generate the plasmid expressing the secreted form of the extracellular domain of zebrafish Tie1 (amino acids 1-747) and Tie2 (amino acids 1-745), DNA fragments encoding extracellular region of *tie1* and *tie2* followed by the Fc region of human immunoglobulin G (amino acids 100-330) and an HA tag were amplified by PCR, and inserted to pCS2+ vector, namely pCS2+-zTie1-Fc-His and pCS2+-zTie2-Fc-His plasmids, respectively. pCS2+-Fc-His plasmid was generated from pCS2+-zTie1-Fc-His plasmid without tie1 sequence after its signal sequence (amino acids 17-747).

A cDNA fragments encoding zebrafish *angpt1*, *angpt2a*, and *angpt2b* without signal sequence were amplified by PCR using cDNA library from 72 hpf zebrafish embryo as a template, then inserted into pCS2+ vector fused with N-terminal preprotrypsin leader sequence and HA tag, namely pCS2+-FLAG-zAngpt1, pCS2+-FLAG-zAngpt2a and pCS2+-FLAG-zAngpt2b plasmids.

pTol flt1:Gal4FF plasmid was constructed by inserting the Gal4FF cDNA into the pTol flt1 vector (Kwon et al., 2013). cDNA fragment encoding zebrafish Foxo1a was amplified by PCR and subcloned into pEGFP-C3 vectors (Takara Bio Inc.) to generate the expression plasmids. Then, EGFP-Foxo1a cDNA was subcloned into the pTol2-fli1 vector (Kwon et al., 2013) to generate the pTol fli1:EGFP-Foxo1a plasmid.

### *In vitro* binding assay

To produce recombinant zTie1-Fc-His, zTie2-Fc-His, Fc-His, FLAG-zAngpt1, FLAG-zAngpt2a, and FLAG-zAngpt2b, 293T cells were transfected with pCS2+-zTie1-Fc-His, pCS2+-zTie2-Fc-His, pCS2+-Fc-His, pCS2+-FLAG-zAngpt1, pCS2+-FLAG-zAngpt2a and pCS2+-FLAG-zAngpt2b plasmids, respectively, using 293 fectin transfection reagent (Gibco), and cultured in DMEM (Wako) supplemented with 10% FBS for 2 days.

Binding of zTie to zAngpt was performed in accordance with the methods of Fukuhara et al. (2008) with some modifications (Fig. 2B). Initially, protein G sepharose beads (GE Healthcare Life Science, Piscataway, NJ) were incubated with supernatants of Fc-His, sTie1-Fc-His or sTie2-Fc-His for 2 hours at 4°C. After washing four times with binding buffer (50 mM Tris-HCl at pH 7.5, 100 mM NaCl, 0.02% Triton X-100), protein bound beads were incubated with supernatants of FLAG-Angpt1, FLAG-Angpt2a or FLAG-Angpt2b for 2 hours at 4°C. After washing with binding buffer four times, the precipitates were subjected to Western blot analysis with anti-penta-His (1:2000, Qiagen) and anti-FLAG M2 (1:4000, Sigma-aldrich) antibodies to quantify the co-precipitated Tie and Angpt proteins, respectively. Proteins reacting with primary antibodies were visualized by the Immobilon forte western HRP substrate (Millipore) for detecting peroxidase-conjugated secondary antibodies and analyzed with an ChemiDoc Touch (Bio-rad).

### Detection of zebrafish Tie phosphorylation

293T cells were cultured until they were 90% confluent. Cells were transfected with pCS2+-zTie1-HA and pCS2+-zTie1KD-HA plasmids using 293 fectin transfection reagent, and cultured in DMEM supplemented with 10% FBS for 9-12 hours. After starvation in 0.5% FBS, DMEM overnight, the cells were stimulated with 500 ng/ml COMP-ANG1 (Cho et al., 2004) for 20 minutes. Cells were lysed in RIPA buffer containing 50 mM Tris-HCl at pH 7.5, 100 mM NaCl, 1% NP-40, 5% Glycerol, 5 mM sodium orthovanadate, 10 mM sodium fluoride, 1 x protease inhibitor cocktail, and 1 mM DTT at 4 °C for 10 minutes, and centrifuged at 15,000 g for 15 minutes at 4 °C. The supernatant was used for Western blot analysis.

To detect the phosphorylated Tie1, aliquots of total cell lysate were subjected to SDS-PAGE and Western blot analysis with anti-phosphotyrosine (1:3000, Cell signaling) antibody. The total contents of Tie1-HA and Tie1KD-HA were assayed in a parallel run using anti-HA antibody (1:2000, Roche). As a loading control, mouse anti-β-actin antibody (1:6000, Sigma-aldrich) was used. Image files were processed and analyzed with ImageJ.

### Fluorescence-activated cell sorting (FACS)

Trunk and tail region from 47 hpf *Tg(kdrl:EGFP);Tg(lyve1:DsRed) tie1*^−/−^ embryos with blood flow and their *tie1*^+/?^ siblings were manually dissected and collected into a low binding 12 well plate. After washed in E3 media, samples were incubated with 1 ml of lysis solution containing collagenase P (Roche) and TrypLE Express Enzyme (Gibco) for 20 minutes under occasional pipetting. Digestion was terminated with 200 μl of stop solution (PBS with 30% FBS and 6 mM calcium chloride). The dissociated cells in suspension medium (phenol red free DMEM (Life Technologies] with 1% FBS, 0.8 mM calcium chloride, 50 U/ml penicillin, and 0.05 mg/ml streptomycin) were subjected to cell sorting using a FACS Aria III cell sorter (BD Bioscience). EGFP and DsRed double positive cells were collected as venous and lymphatic endothelial cells for further RNA preparation.

### RNA sequencing

Total RNA was prepared from *kdrl*:EGFP^+^/*lyve1*:DsRed^+^ ECs of *tie1*^−/−^ embryos and their *tie1*^+/?^ siblings using the NucleoSpin XS kit (Macherey-Nagel, 740902.50) according to the manufacturer’s instructions. Reverse transcription and cDNA library preparation were performed with a SMART-Seq v4 Ultra Low Input RNA Kit for Sequencing (Clontech, 634888). cDNA was fragmented with a Covaris S 220 instrument (Covaris). Libraries for RNA-Seq were prepared using a NEBNext Ultra II Directional RNA Library Prep Kit for Illumina (New England Biolabs, E7760) and sequenced on the NextSeq500 (Illumina) as 75 bp single-end reads. RNA-Seq data were trimmed using Trim Galore version 0.6.6 (https://www.bioinformatics.babraham.ac.uk/projects/trim_galore/) and Cutadapt version 2.8 (Martin, 2011). The quality of reads was checked and filtered using FastQC version 0.11.9 (http://www.bioinformatics.babraham.ac.uk/projects/fastqc). The reads were mapped to a reference genome GRCz11 using HISAT2 version 2.2.1 (Kim et al., 2019), and the resulting aligned reads were sorted and indexed using SAMtools version 1.7 (Li et al., 2009). Relative abundances of genes were measured in TPM using StringTie version 2.1.4 (Kovaka et al., 2019; Pertea et al., 2015). Plots were created in Python using the pandas, matplotlib, NumPy, and seaborn libraries. Differentially expressed genes (DEGs) were identified using TCC-GUI (Su et al., 2019; Sun et al., 2013). Genes with q-value < 0.05 (5% false discovery rate, FDR) were considered to be differentially expressed. To identify the GO biological process, enrichment analysis was performed on the list of 1,444 differentially expresses genes using Metascape v.3.5 (Zhou et al., 2019).

### Data analysis and statistics

Data were analyzed using GraphPad Prism software or Excel and were presented as mean ± s. d. Sample numbers were indicated in figure legends. The statistical significance of two groups was determined using Student’s t-test or Welch’s t-test. In case of Figure 7B, a Mann-Whitney test was performed instead.

## ACKNOWLEDGEMENT

We thank Kenta Terai, Takefumi Kondo, and Yukari Sando (NGS core facility of the Graduate Schools of Biostudies, Kyoto University) for supporting the RNA-Seq analysis. We are grateful to B. Hogan (The University of Melbourne) and K. Koltowska (Uppsala University) for the plasmid for *Tg(10×UAS:Vegfc)*, K. Koltowska (Uppsala University) for the detailed protocol for Prox1 immunostaining, K. Kawakami (National Institute of Genetics) for the Tol2 system, G.Y. Koh (Korea Advanced Institute of Science and Technology (KAIST)) for COMP-ANG1, M. Sone, T. Satoh, K. Hiratomi, T. Babazono and E. Hanimura for technical assistance, K. Shioya and K. Konno for fish care.

## AUTHOR CONTRIBUTIONS

Conceptualization: N. Morooka, N. Mochizuki, H.N.; Methodology: N. Morooka, N.G., M.F. M.H.; Software: N. Morooka; Validation: N. Morooka, N.G., K.S., M.H., H.N.; Formal analysis: N. Morooka, N.G., H.N.; Resources: K.A., S.S.-M.; Data Curation: N. Morooka, H.N.; Writing - original draft: N. Morooka, H.N.; Writing - review & editing: S.S.-M., H.N.; Visualization: N. Morooka, H.N.; Supervision: S.S.-M., N. Mochizuki; Project administration: N. Mochizuki, H.N.

## DATA AVAILABILITY

RNA-seq data were deposited in the Gene Expression Omnibus under accession number GSE240329. The reviewers can access our private data using the following links: https://www.ncbi.nlm.nih.gov/geo/query/acc.cgi?acc=GSE240329. The data will become public after publication of our paper.

## MOVIES

**Movie 1. Normal secondary sprouting in a wild-type (WT) embryo.**

Time-lapse recording in the trunk of a *Tg(kdrl:EGFP);Tg(lyve1:DsRed)* WT embryo from 36 hpf taken every 30 min by confocal microscopy. This Tg embryo expresses EGFP (green) in all endothelial cells (ECs) and DsRed (magenta) in venous and lymphatic ECs. Elapsed time (h:min:sec). Lateral view, anterior to the left. Secondary sprouts from the posterior cardinal vein (PCV) migrate to the horizontal myoseptum to form the parachordal lymphangioblasts (PLs), a pool of lymphatic precursors.

**Movie 2. Disrupted secondary sprouting in a *tie1* mutant embryo.**

Time-lapse recording in the trunk of a *Tg(kdrl:EGFP);Tg(lyve1:DsRed) tie1^−/−^* embryo from 37 hpf taken every 30 min by confocal microscopy. Elapsed time (h:min:sec). Lateral view, anterior to the left. In the *tie1* mutant, sprouting ECs extend their protrusions but do not migrate dorsally.

